# DegCre: Probabilistic association of differential gene expression with regulatory regions

**DOI:** 10.1101/2023.10.04.560923

**Authors:** Brian S. Roberts, Gregory M. Cooper, Richard M. Myers

## Abstract

Differential gene expression in response to perturbations is mediated at least in part by changes in binding of transcription factors (TFs) and other proteins at specific genomic regions. Association of these cis-regulatory elements (CREs) with their target genes is a challenging task that is essential to address many biological and mechanistic questions. Many current approaches rely on chromatin conformation capture techniques that identify spatial proximity between genomic sites to establish CRE-to-gene associations. These methods can be effective but have limitations, including resolution, minimal detectable interaction distance, and cost. As an alternative, we have developed DegCre, a non-parametric method that evaluates correlations between measurements of perturbation-induced differential gene expression and differential regulatory signal at CREs to score possible CRE-to-gene associations. It has several unique features, including the ability to: use any type of CRE activity measurement; yield probabilistic scores for CRE-to-gene pairs; and assess CRE-to-gene pairings across a wide range of sequence distances. We apply DegCre to three data sets, each employing different perturbations and containing a variety of regulatory signal measurements, including chromatin openness, histone modifications, and TF occupancy. To test their efficacy, we compare DegCre associations to HiC loop calls and to CRISPR validated interactions, with both yielding good agreement. We demonstrate the identification of perturbation direct target genes with DegCre confirm the results with previous reports. DegCre is a novel approach to the association of CREs to genes from a perturbation-differential perspective, with strengths that are complementary to existing approaches and allow for new insights into gene regulation.

## Introduction

The regulation of gene expression occurs through the interaction of transcription factors (TFs) and other proteins with genomic regions, or cis-regulatory elements (CREs) (The ENCODE Project Consortium et al. 2020). Because CREs can act at considerable distances away from any given target gene, in some cases skipping over intervening genes, matching CREs to their target genes is challenging (Rao et al. 2014; Song et al. 2019; Dixon et al. 2012; Javierre et al. 2016; Schoenfelder and Fraser 2019). Such knowledge, however, is of high value for both basic and applied biology. Examples include better interpretation of how genetic variation in a CRE leads to molecular and phenotypic effect (Ulirsch et al. 2016; van Arensbergen et al. 2019; Song et al. 2019; Nasser et al. 2021) and better prediction of the effects of perturbations, such as drug treatment, on gene expression levels in a cell or tissue (Thormann et al. 2018; Carleton et al. 2020; Cholico et al. 2022).

Promoters are CREs very near to the transcription start site (TSS) of a gene and are readily associated with that gene’s expression by the basic principles of transcription initiation (Haberle and Stark 2018). As sequence distance from the TSS increases, the association of CREs, such as enhancers, with a given gene’s regulation becomes increasingly uncertain. Analysts often place a threshold on TSS to CRE distances under which there is putatively high confidence for the association. These thresholds range from ∼1 kilobase (kb) to 100s of kb, with little justification provided for any given choice (Kamal et al. 2023; You et al. 2021; McDaniel et al. 2016; Wang et al. 2015). It is unlikely that any single TSS to gene distance threshold is appropriate in all contexts, and categorical thresholds generally result in limitations that impact all downstream analyses.

The maturation of chromatin conformation capture technology such as HiC has enabled the association of genomically distal CREs to genes through their spatial proximity in nuclei to target-gene promoters (Rao et al. 2014; Javierre et al. 2016; Kloetgen et al. 2020; Xu et al. 2022; Meng et al. 2023). The high resource usage is one of the key limitations of these approaches. Moreover, the majority of HiC associations, or “loops”, span distances greater than ∼50 kb, limiting their utility to identify CRE-to-gene associations spanning shorter distances. Undoubtedly, CREs closer to TSSs harbor considerable regulatory activity; yet it is also unlikely that all CREs in this distance range are in fact relevant to the regulation of that gene, especially in response to a targeted perturbation.

Previous work by others has demonstrated approaches to CRE-to-gene association that do not require chromatin conformation data. Many of these approaches are based on the observation that gene transcription levels and measurements of CRE regulatory activity are correlated (Ernst et al. 2011; Sheffield et al. 2013; He et al. 2014; Cao et al. 2017; Li et al. 2019). Generally, these methods generate models of CRE activity and gene expression within a given cell type across the genome, yielding predictions of associations. Validation of these methods by comparison to chromatin capture data yields good agreement in most cases. One possible limitation of these methods is that many require supervised training which could lead to underperformance when applied to untrained contexts. Also, these methods are tailored to specific data types, lacking flexibility in inputs. Nevertheless, this work has established the ability of correlational analysis to identify CRE-to-gene associations.

Studies that enable the association of CREs to genes, whether through HiC approaches, correlational methods, or other techniques, often focus on static conditions, generating data of CRE activity and gene expression in a single context, with notable exceptions (Adamson et al. 2016; Reed et al. 2022; Xu et al. 2022). In contrast, a differential system in which two or more conditions are compared may allow for a new conceptual approach to CRE-to-gene association. This approach correlates gene expression and CRE activity not across genes and CREs in a single context, but between the same gene and same CRE across two or more conditions. Correlations in CRE activity between conditions that are concordant with gene expression changes may provide evidence of their association. Such associations would be dependent on the variable factor between the input conditions, perhaps limiting their generality, but also would be more informative of the variable effects and potentially lead to insights into the gene regulatory mechanisms at work.

We present a method, DegCre, that probabilistically associates CREs to target gene TSSs over a wide range of genomic distances. The premise of DegCre is that true CRE to differentially expressed gene (DEG) pairs should change in concert with one another as a result of a perturbation, such as a drug treatment or differentiation protocol. DegCre is a non-parametric method that estimates an association probability for each possible pair of differential CRE and DEG. It considers CRE-DEG distance but avoids arbitrary thresholds. Because DegCre uses rank-order statistics, it can use various types of CRE-associated data, including DNase hypersensitivity, ATAC-seq, and ChIP-seq against either histone marks or TFs. It produces significant associations with a wide range of TSS to CRE distances, up to an upper limit subject only to computational burden, with 1 Mb being quite feasible.

We apply DegCre to three distinct collections of data, including cells subject to drug treatment and differentiation protocols. We compare DegCre associations to HiC loops, CRISPR validated links, and Activity By Contact (ABC) scores (Nasser et al. 2021) that employ both HiC and regulatory signal data. While DegCre makes CRE-to-gene predictions well below the limit of genomic distance detection of HiC data, we find excellent agreement among areas of predictive overlap. We also show the complementary value of DegCre’s scoring metric, which provides, for example, the ability to sum probabilities for all candidate CREs for a given gene to infer the total number of CREs regulating that gene. We use this approach to identify direct targets of a perturbation by identifying genes regulated by numerous CREs.

## Results

### DegCre operation and algorithm

The name “DegCre” was chosen to reflect that, instead of binary associations, DegCre produces association probabilities, or the “degree” to which the DEG to CRE association is likely to be true. For the purposes of this discussion, we define a differentially expressed gene (DEG) as a gene whose expression is potentially different between two contexts, such as before and after drug treatment or differentiation. The term “significant DEG” will mean DEGs for whom the significance of a statistical measure of expression change surpasses a defined threshold (α). The term “CRE” (cis-regulatory element) will be used to denote a genomic region with regulatory signal above background at a chosen level. Such regulatory signals can include chromatin openness, transcription factor (TF) occupancy, and histone post-translational modifications, amongst others. We understand that robustly establishing that any given genomic region is truly a CRE requires additional lines of evidence, such as targeted inhibition and activation, but will use CRE in this study to refer to any region exhibiting condition-differential signal for the selected type of CRE-measurement.

We intend for users to apply DegCre to experimental designs which consider multiple states, such as response to a perturbation, and include measurements of gene expression and regulatory signal at CREs (Figure 1A). DegCre takes in differential gene expression measurements, defined as the p-value from comparing expression levels between conditions, along with the transcription start sites (TSSs) of those genes. DegCre also requires p-values of differential signal at CRE regions from the same conditions, such as can be generated by methods like csaw (Lun and Smyth 2016). Optimally, DegCre also uses the fold-changes of both DEG and CRE signal to measure effect direction concordance on the assumption that CRE activity should correlate with increased expression and vice versa, although this is not required and for some experimental designs may not be desired.

**Figure 1.**
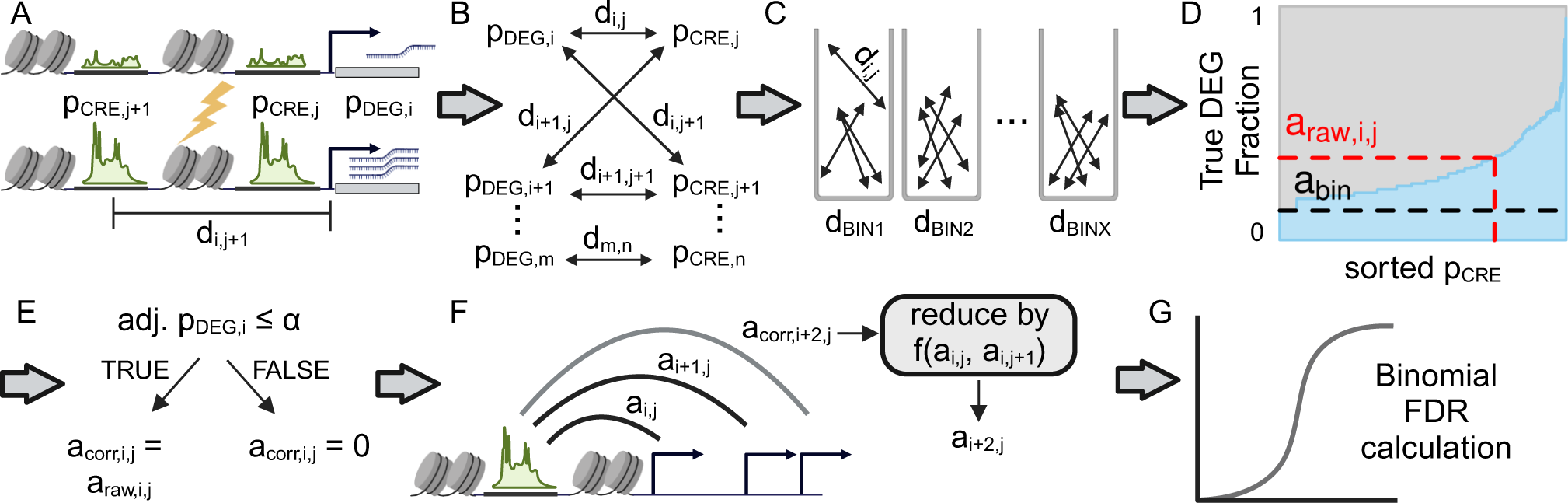
Graphical overview of the DegCre algorithm. A. DegCre requires as inputs differential p-values for CRE signal and differential gene expression (p_CRE_ and p_DEG_). DegCre also needs genomic distances between CREs and TSSs (d) as input. The lightning bolt indicates a perturbation has occurred to the lower depiction. B. DegCre creates also possible associations between each CRE and TSS within a specified maximum distance. C. DegCre bins associations by their distance, d, according to a heuristic that balances resolution versus maintaining the p_CRE_ distribution (Methods). D. DegCre calculates a raw association probability, a_raw,i,j_, for a given p_CRE,j_ by finding the fraction of expected true DEGs in the set of associations in the same distance bin and with a p_CRE_ equal to or less than (more significant) as p_CRE,i_. Plot shows actual data from ATAC-seq at 120 minutes from Reed et al. E. DegCre corrects the raw association probability if the association does not involve a true DEG. F. For CREs with multiple associations (nearly all CREs) associations across larger genomic distances are penalized by the probabilities that the CRE is associated to nearer DEGs. G. The false discovery rate (FDR) of the association is calculated based on a binomial distribution that uses the bin null association probability, a_bin_, as the success probability. Created with BioRender.com.

The DegCre algorithm begins by defining all pairwise associations between DEGs and CREs and calculating the genomic distance between each CRE and the TSSs of each DEG (Figure 1A and B, Methods). A DegCre association is thus based on three measurements: a DEG p-value, a CRE p-value, and a genomic distance. DegCre considers only possible associations within a given maximum distance, which is by default set to be 1 Mb. Specification of large maximum distances is possible but increases the number of considered associations and computational burden.

DegCre next bins associations by distance (Figure 1C, Methods). The bin sizes are defined for each experiment such that they contain an equal number of associations, (Methods). The goal of this step is to define bins such that each CRE can be compared against other CREs at similar distances to yield scores for individual CRE-DEG associations that can be compared to the average distance-normalized association; true CRE-DEG pairs should have higher signal than randomly selected CRE-DEG pairs with similar distances. To select the number of associations per bin, DegCre employs a heuristic that attempts to balance resolution (smaller bins with fewer associations) against the robustness of each bin, which is a function of the similarity of the CRE p-value distribution in each bin compared to the global CRE p-value distribution (Methods). It is important to maintain uniformity of the bin-wise CRE p-value distribution because subsequent calculations compare the CRE p-value for any given association to the distribution of all CRE p-values of its distance bin. Thus, if bins are too small, and harbor only highly significant or only insignificant p-values, then true pairs cannot be distinguished from the background. In contrast, if bins are too large, the distance normalization becomes irrelevant as distances with biologically very different priors (in terms of likelihood for a random association to represent a real association) are conflated.

For each bin, DegCre calculates raw association probabilities, represented by “a” in figures and formulas to avoid confusion with significance probabilities (Figure 1D). For a given association with a given CRE p-value, DegCre finds all associations within the same bin that have a CRE p-value (in the same effect direction if available) with equal or greater significance (Methods). DegCre then calculates the expected number of true DEGs within that set of associations and divides it by the set size to obtain a true DEG fraction (Figure 1D, Methods). We call this value the raw association probability, a_raw_, an estimation of the probability the association connects a CRE to the change in expression of a target gene. This calculation a_raw_ is determined by association distance, CRE p-value, and DEG p-value, and makes no assumptions of underlying distributions of those inputs.

Although there is a unique a_raw_ value for each unique CRE p-value, a_raw_ values essentially represent a set-level probability, as it reflects the cumulative probability of all associations with that degree of CRE significance or greater. DegCre therefore corrects a_raw_ by the probability that an association involves a significant DEG to generate a_corr_ (Figure 1E). DegCre applies this correction to associations with adjusted DEG p-values passing a selected α (Methods). The choice of alpha can be guided by an included optimization function. A given CRE will generally have associations to multiple significant DEGs. All else being equal, we assume that the association most likely to be real is to the nearest DEG. DegCre thus considers all significant DEG associations for a given CRE and reduces a given a_corr_ weighted by the a_corr_ values of associations containing that CRE that have shorter association distances (Methods), producing the final reported association probability (Figure 1F).

For FDR estimation of the association probability, DegCre considers the raw association probability of each bin without regard to CRE p-value, a_bin_ (Figure 1D). The a_bin_ value depends on distance-based binning only (i.e., it reflects the scores derived from all possible CREs within a given distance range) and we consider it to be a suitable null hypothesis for the effect of CRE p-values on association probabilities. Accordingly, DegCre calculates an association probability FDR based on the binomial cumulative distribution function using a_bin_ as the trial success probability (Figure 1G; Methods).

DegCre is implemented as an open-source R package (Methods). It operates within the convenience of GenomicRanges (Lawrence et al. 2013) framework. The generated outputs enable manipulation with existing operations. It includes functions for secondary calculations, visualization of results, and conversion of the results into other formats.

### Data sets for the demonstration of DegCre analysis

DegCre operates on measurements of gene expression and signal at CREs in response to a perturbation to generate association probabilities between genes and CREs. We therefore sought to test DegCre using datasets from perturbation experiments such as those done using drug treatment and differentiation protocols.

First, we analyzed data published by McDowell et al. derived from the treatment of A549 cells by dexamethasone (McDowell et al. 2018). Dexamethasone is a specific and potent agonist of the glucocorticoid receptor (NR3C1), a well-studied nuclear receptor (Vettorazzi et al. 2022) that is known to change expression of many target genes upon stimulation. This data set includes an extensive time course as well as measurements of RNA, DNase hypersensitivity, chromatin immunoprecipitation sequencing (ChIP-seq) of mono- and trimethylation of histone H3 at lysine 4 (H3K4me1 and H3K4me3) and acetylation of histone H3 at lysine 27 (H3K27ac), and ChIP-seq of several transcription factors and co-activators including NR3C1 itself.

We used a second data set generated in our lab involving the activation of a different nuclear receptor (Savic et al. 2015). Specifically, Savic et al treated HT-29 cells with rosiglitazone, an antidiabetic drug acting through activation of peroxisome proliferator-activated receptor gamma (PPARG). PPARG is a nuclear receptor that upon activation forms heterodimers with retinoid X receptors (RXRs) and translocates to the nucleus, modulating gene expression changes (Han et al. 2017). This study includes RNA-seq, H3K27ac ChIP-seq, and RNA Polymerase II (RNA Pol 2) ChIP-seq at 24 and 48 hours after treatment.

A third dataset we used involved another perturbation with direct effects on gene regulation: the stimulation of immune cells with chemokines. Interferon gamma (IFNγ) is a powerful chemokine resulting in the phosphorylation and consequent activation of the transcription factor STAT1 (Schneider et al. 2014). Reed et al stimulated THP-1 monocytes with IFNγ and lipopolysaccharide and collected samples at eight timepoints (Reed et al. 2022). Measurements included in this data set are RNA-seq, H3K27ac ChIP-seq, and ATAC-seq data. This study also generated genome-wide HiC measurements of chromatin conformation. Because evidence of spatial proximity between promoters and distal CREs provides an experimental inference of CRE-DEG pairs, these data allow for comparisons of DegCre associations to an orthogonal assay.

These data sets are publicly available and complete accession details are provided in Supplemental File 1. For ChIP-seq, ATAC-seq, and DNAse hypersensitivity data we aligned raw reads when necessary or used author-supplied BAM files. We then applied the R package csaw (Lun and Smyth 2016) to generate regions of differential signal as GRanges objects (Methods). For RNA-seq, we used author-provided counts and determined differential expression between treatments and timepoints using DESeq2 (Methods). We associated the differential expression results with gene TSSs as defined in the EPDnew database (Meylan et al. 2020; Dreos et al. 2015) and created GRanges objects (Methods).

### Characteristics of DegCre associations

We applied DegCre analysis to each of the three datasets described above, which take at most a few minutes on a regular desktop computer using default settings, including measuring CRE-to-promoter pairings up to 1 Mb of distance (Methods). We provide the outputs as text files in Supplemental File 2. The total number of associations passing an FDR threshold (by default set to 0.05) varies across each dataset from hundreds to tens of thousands (Supplemental Figure 1A). The total number of significant DEGs largely determines the number of significant DegCre associations (Supplemental Table 1, Methods); that is, experiments with more DEGs yield more DEG-CRE pairs (Supplemental Figure 1B), because only associations involving a significant DEG can potentially pass reasonable FDR thresholds (Methods). The total number of CREs with nominally significant differential signal p-values has less effect on the total number of significant DegCre associations (Supplemental Figure 1C).

The number of significant associations generally decreases with increasing genomic distance between DEG TSSs and CREs (Figure 2A-C, Supplemental Figures 2-6). While this likely reflects real biology, this distance dependence also reflects the fact that DegCre weights equivalently significant associations for a given single CRE in favor of the ones spanning shorter genomic distance (Methods). Thus, longer associations are likely to pass FDR correction only if there are no other shorter associations of equal or greater strength involving that CRE. Associations from some data types from McDowell et al. display a slower decrease versus distance compared to others (Supplemental Figure 3). For example, associations from ChIP-seq from H3K4me3 show a much steeper decrease compared to those from H3K4me1 or EP300. This observation is consistent with the known binding profiles of these factors, as H3K4me3 is primarily associated with promoters and promoter-proximal CREs, while H3K4me1 and EP300 are primarily associated with enhancers and other distal CREs (The ENCODE Project Consortium et al. 2020; Visel et al. 2009).

**Figure 2.**
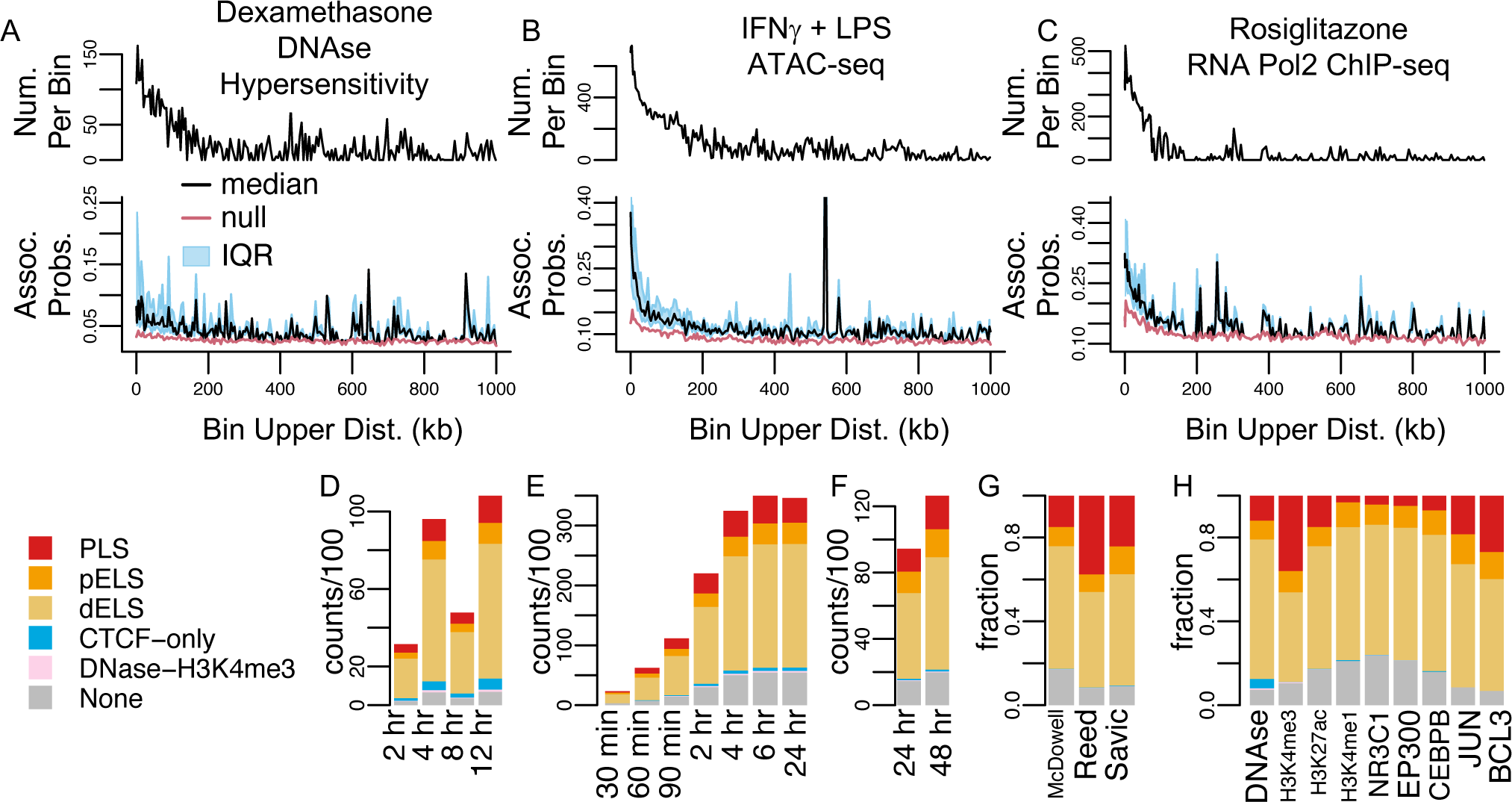
Characteristics of DegCre associations. A-C. The black line in the upper plot half displays the number of DegCre associations per bin that pass the indicated FDR. The bottom half displays the per bin DegCre association probability. The common x-axis shows for each bin the association distance from TSS to CRE. Each bin comprises a range of distances with the upper bound of that range plotted here. The black line indicates the median value for each bin and the blue region indicates the interquartile range (IQR). The red line shows the per bin probability considering only the bin distance, used as the null in the DegCre FDR calculation (Methods). DegCre associations are shown from: A.) DNase hypersensitivity data at eight hours from McDowell et al., B.) ATAC-seq data at two hours from Reed et al., C.) RNA Pol2 ChIP-seq data at 24 hours from Savic et al. D-H. Bars show the counts or fraction of differential regions with a significant (FDR less than 0.05) DegCRE association overlap ENCODE cCRE annotations. Plots are based on: D.) DNase hypersensitivity data from McDowell et al., E.) ATAC-seq data from Reed et al., F.) RNA Pol2 ChIP-seq data hours from Savic et al. (same data as A-C). G. Fractions are shown for H3K27ac ChIP-seq data from McDowell et al. at eight hours, Reed et al. at 2 hours, and Savic et al. at 24 hours. H. Fractions are from McDowell et al. at eight hours for the indicated data types. Abbreviations for ENCODE cCREs are: PLS, promoter like sequence; pELS, proximal enhancer-like sequence; dELS, distal enhancer like sequence.

The values of the association probabilities passing the FDR threshold vary from just above the null probability for a given bin (red lines in bottom halves of Figures 2A-C, Supplemental Figures 2-6) to higher values closer to 100% (interquartile ranges shown in bottom halves of Figures 2A-C, Supplemental Figures 2-6). These ranges illustrate a key characteristic of DegCre associations. The DegCre FDR estimates whether the association probability exceeds the null, bin-wide probability due to the input of the differential CRE signal. However, those associations that pass a chosen FDR threshold can still be further stratified by the association probability itself. For example, we infer, to a reasonable approximation, that a DegCre association with a probability of 0.9 will be three times more likely to confirm in an orthogonal assay to one with 0.3, with both passing a chosen FDR threshold.

The DegCre algorithm uses differential CRE signal in a non-parametric, rank-based analysis without thresholding (Methods). DegCre is agnostic to the methods of assigning differential CRE significance p-values to genomic regions. We used csaw with defined windows (20 bp for TFs, 200 bp for histone marks and open chromatin signals) spanning hg38 to assess the level of differential signal between conditions for each type of CRE measurement. We then calculated csaw p-values for the windows passing signal thresholds prior to DegCre analysis (Methods). We did not use peak calling for any CRE dataset. A possible concern with this processing strategy is that it could lead to DegCre associations involving CREs of dubious regulatory potential. Accordingly, for all datasets we intersected ENCODE-defined cCRE annotations (The ENCODE Project Consortium et al. 2020) with all CREs predicted by DegCre to target a DEG with an FDR less than 0.05 (Figure 2A-H, Supplemental Figure 7). More than 80% of these DegCre associations overlapped an ENCODE cCRE annotation, with many data sets having higher overlap rates (Supplemental Figure 7). Most CREs annotate as proximal or distal enhancer-like sequences (pELS and dELS), consistent with these being the most prevalent annotations in the ENCODE cCRE set. We looked at H3K27ac ChIP-seq across all three data sets because it was the only CRE signal that was present in all three. We found that CREs in significant DegCre associations from Reed et al. and Savic et al. overlapped promoter-like signature (PLS) more frequently than did those from McDowell et al. (Figure 2H). Given that the perturbation in McDowell et al was targeted to NR3C1, this observation is expected given NR3C1’s known bias towards distal CREs in response to activation (Thormann et al. 2018; Vettorazzi et al. 2022).

We also note that the total number of significant DegCre associations either consistently increases or increases and plateaus with time after perturbation for all three datasets. However, the relative fraction of ENCODE annotations tended to remain stable over time (Figure 2D-H, Supplemental Figures 1 and 7). This time dependence may arise from the accumulation of secondary gene regulation that occurs post-perturbation. Regardless, the time elapsed from the perturbation does not appear to affect the types of ENCODE cCREs involved in DegCre associations.

We next examined the frequency of ENCODE cCRE annotations for the nine CRE-associated data types in the McDowell et al. study (Figure 2H). For H3K4me3 ChIP-seq, we observed an enrichment for PLS annotations, consistent with the known prevalence of this mark at active promoters, and we also saw promoter enrichment for JUN and BCL3, again as expected. In contrast, we saw enhancer enrichment for NR3C1 and EP300 ChIP-seq CREs, as expected for the CREs associated with these marks. Thus, DegCre’s inferences are reflective of the nature of the input data, yielding differing associations from differing input data types.

### Comparison of DegCre associations to HiC loops

We sought to compare DegCre associations to HiC-derived loop calls from Reed et al., who generated HiC loop calls using Sip (Rowley et al. 2020) at an FDR < 0.05 (Reed et al. 2022). We used liftover (Lawrence et al. 2009) to convert these loops to hg38 coordinates (Methods).

We excluded 18 hg38 loops with loop sizes (defined as genomic distance from anchor-to-anchor midpoints) smaller than 30 kb after liftover because all loop sizes were 30 kb or greater prior to liftover. The remaining loops spanned up to 28,617,181 bp with a median size of 390,000 bp (Supplemental Figure 8A). Significant (FDR <=0.05) DegCre association distances (from CRE to gene TSS) derived from both ATAC-seq and H3K27Ac ChIP-seq data from Reed et al. are mostly shorter than HiC loops, especially at time points past 30 minutes (Supplemental Figure 8B-D), occurring at distances down to zero (i.e., at the TSS itself). Also, we set the maximum DegCre association distance to 1 Mb due to the increased computational burden at higher maximum distances. Accordingly, to evaluate the overlap of these two measures of distal CRE associations to DEGs, we considered only DegCre associations and HiC loops in the distance range of 30 kb to 1 Mb.

By design, significant DegCre associations can occur only between significantly expressed DEGs and CREs, as marked by the input assay (here ATAC-seq and H3K27Ac ChIP-seq, see Methods); that is, if a gene is not differentially expressed between conditions, or if a CRE does not show differential activity, there will be no DegCre association prediction. To compare DegCre associations to HiC loops, we thus considered only HiC loops with one anchor overlapping a CRE with a significant DegCre association. For both ATAC-seq and H3K27Ac ChIP-seq, approximately one-third of the HiC loops met this criterion (Supplemental Figure 9, compare A and B). For HiC loops that do overlap a DegCre association CRE, we placed each HiC loop into one of four categories: (1) it overlaps the TSS of the same gene that the DegCre association does (Figure 3A), (2) it overlaps the TSS of a different gene than the DegCre association and that different gene is also significantly differentially expressed (Figure 3B), (3) it overlaps the TSS of a different gene than the DegCre association and that different gene is *not* significantly differentially expressed, (4) it does not overlap the TSS of a gene. We found that most HiC loops fall into the last two categories (Supplemental Figure 9, compare C and D). These loops are difficult to interpret in comparison to DegCre associations. They associate CREs to regions that DegCre does not consider, including sites that are not TSSs (such as distal CREs) or to TSSs of non-differentially expressed genes. Either category cannot be compared to DegCre associations. Accordingly, we considered only the first two categories further.

**Figure 3.**
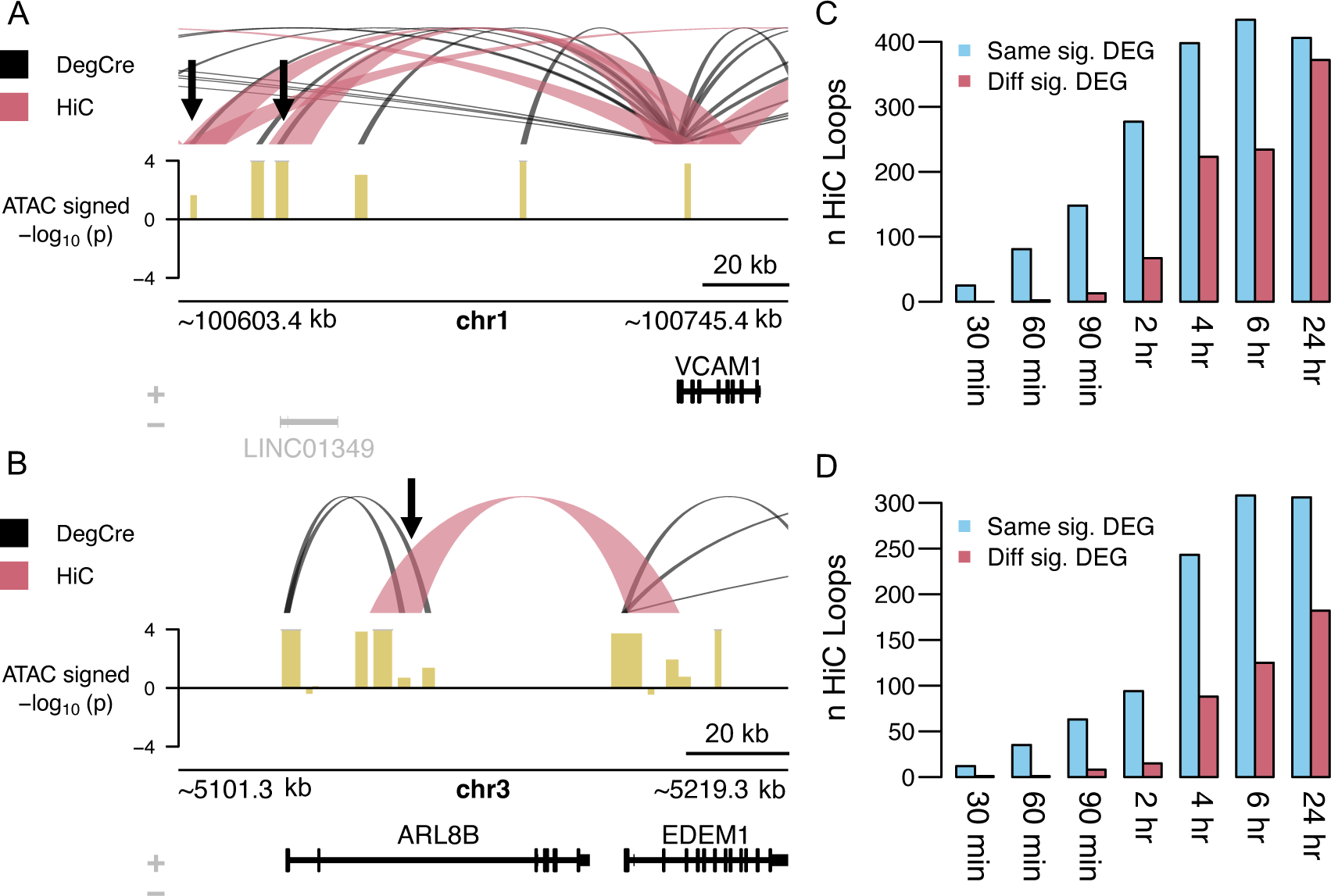
Comparison of DegCre associations to HiC loops. A. For ATAC-seq data from Reed et al at the 120-minute timepoint, DegCre associations with an FDR less than 0.05 and an association distance greater than 20 kb are shown in black. HiC loops with an FDR less than 0.05 and a loop distance less than 1 Mb are shown in light red. Gene names in black indicate significant differential expression. Black arrows indicate distal CREs that both DegCre and HiC link to the VCAM1 TSS. The signal track (yellow) shows the −log_10_ of the differential ATAC signal multiplied by the sign of the log fold-change. B. Same plotting conventions as A but the black arrow indicates a group of CREs for for which DegCre and HiC assign to the TSSs of different, significant DEGs. C. For ATAC-seq data, the blue bars indicate the number of HiC loops that have one anchor in a CRE with a significant (FDR less than 0.05) DegCre association that link to the TSS of the same DEG as the DegCre association. Red bars indicate the number of HiC loops that have one anchor in a CRE with a significant DegCre association that link to the TSS of the different DEG as the DegCre association. D. Same plotting conventions as C but for H3K27Ac ChIP-seq data.

Comparing the partitioning of the HiC loops into these first two categories across all timepoints and both CRE-defining data types, ATAC-seq and H3K27Ac ChIP-seq, we found that more DegCre associations and HiC loops associate with the same significant DEG than with different significant DEGs (Figure 3C and D). This effect is greater at earlier time points, likely indicating the higher ability of DegCre to properly assign distal CREs to DEGs when considering data temporally closer to the driving perturbation.

### DegCre performance on CRISPR validated associations

A powerful technique for evaluating genomic regions’ potential to regulate genes is through editing or repressing the activity of these regions with CRISPR-Cas9 sequence alteration or CRISPR interference (CRISPRi) (Gasperini et al. 2019; Nuñez et al. 2021). To that end, we obtained a compilation of CRISPR and CRISPRi perturbations of genomic loci and corresponding gene expression measurements from Nasser et al. (Nasser et al. 2021). After liftover to hg38 coordinates, there were 5,685 gene to distal CRE (greater than 500 bp) associations, involving 1,913 unique regions.

Again, DegCre aims to find associations between genomic regions and DEGs with significant changes in response to a perturbation. Accordingly, we filtered the DegCre associations from McDowell et al., Reed et al., and Savic et al. to those containing a significant DEG. However, much of the Nasser et al. data set includes associations that do not validate by CRISPR. While we expect that high probability, low FDR DegCre associations will be CRISPR-validated, we also expect low probability, high FDR DegCre associations to be enriched in the associations lacking CRISPR confirmation. Thus, for these comparisons we retained all DegCre predictions, at any FDR level.

Nasser et al. classified each CRE-to-gene association as “Regulated” or “not” based upon the CRISPR data. We used this binary value in our subsequent analyses. We overlapped the filtered DegCre associations to the CRISPR association data, finding various degrees of overlap for each data set and type (Supplemental File 3). Associations derived from the McDowell et al. data set had generally low levels of overlap with the CRISPR data, making a rigorous comparison impossible. The genes regulated by dexamethasone in this experiment appear to differ greatly from those measured in the Nasser et al. data.

DegCre associations from Reed et al. and Savic et al. had higher overlap rates with the CRISPR data than those from McDowell et al. (Supplemental File 3). We examined the distribution of DegCre association probabilities between those associations designated as “Regulated” versus “not” and found extremely significant differences in several of the distributions by Wilcoxon rank-sum test (p-values from 1.3 x 10^-6^ to < 2.2 x 10^-16^, Figure 4, Supplemental File 3). These results demonstrate agreement between DegCre and CRISPR-based validation. Within the subsets of DegCre associations that overlap the CRISPR validated regions, those from Reed et al. had much broader dynamic range of association probabilities compared to Savic et al. associations (Supplemental Figure 10).

**Figure 4.**
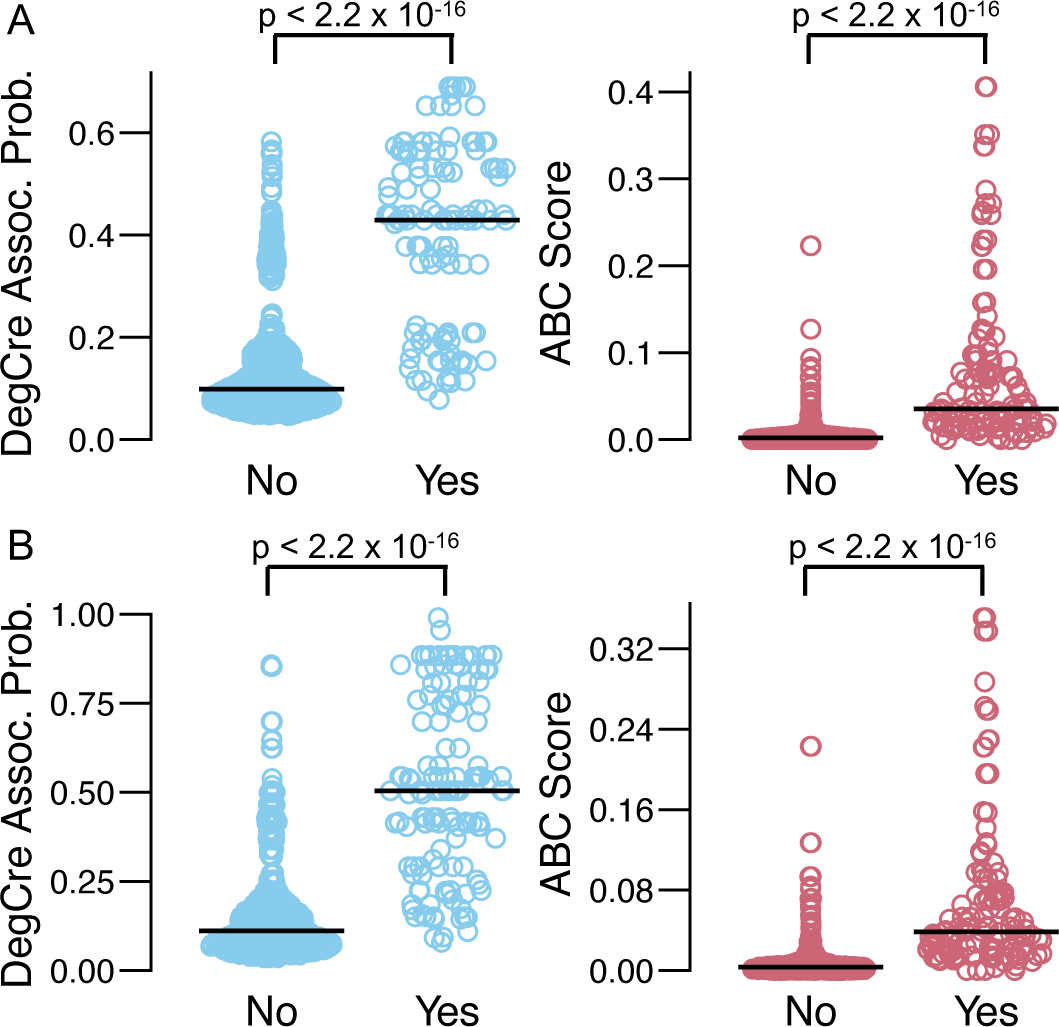
DegCre and ABC association values compared to CRISPR validated enhancer-gene pairs. The blue data points represent DegCre association probabilities based on data at 24 hours from Reed et al. The red circles are ABC scores for the same associations. The data are grouped by whether the association showed regulation of the target gene (“Yes”) or not (“No”) upon CRISPR targeting as given by Nasser et al. The black lines represent the median value. The jitter width of the points is proportional to the local density. The DegCre values are based on A). ATAC-seq data at 1,539 regions and B). H3K27ac ChIP-seq data at 875 regions. The ABC distributions appear very similar between A and B and do share many points but are distinct.

The Nasser et al. data also includes ABC scores for each experimentally tested association. These scores are based on HiC and regulatory signal data. We compared the ABC scores for the subset of CRISPR associations overlapping each DegCre set (Figure 4, Supplemental File 3). The distributions of ABC scores between “Regulated” or “not” associations in these subsets are significantly different in all cases (Figure 4, Supplemental File 3). Many of the ABC Wilcoxon p-values are more significant than the corresponding DegCre p-values. However, many of the Reed et al. DegCre associations compare well to ABC, likely due to the broader dynamic range of DegCre association probabilities present in this set for these regions (Figure 4, Supplemental Figure 10, Supplemental File 3).

### Prioritization of perturbation target genes with DegCre

A key goal of perturbation studies is often to identify the direct target genes among a set of significant DEGs. For perturbations that involve the activation of transcription factors, such as the introduction of ligands to nuclear receptors, “direct targets” means those genes with expression changes that are mechanistically attributable to a change in the genomic occupancy of the nuclear receptor at a CRE that regulates that gene. Identification of direct target genes is desirable because it can lead to insight into the perturbation’s mechanism of action, and it may increase the extensibility of the observed experimental results to other systems.

As DegCre outputs probabilistic scores, it can calculate the expected number of truly associated CREs for each DEG. For a given gene, this is simply the sum of its association probabilities passing a specified FDR cutoff, resulting in an estimation of the number of associated differential CREs that would pass definitive validation (e.g., for a DEG with 10 FDR-passing associations averaging 20% association probability, it would be expected that there are two true CRE-DEG pairs). Thus, genes with many expected associations are more likely to be confidently linked to a driving regulatory signal. We calculated the expected associations per significant DEG for all the tested data sets (Figure 5A, Supplemental Figure 11). The distributions of expected associations per DEG generally showed a slight increase with time followed by a plateau or decrease.

**Figure 5.**
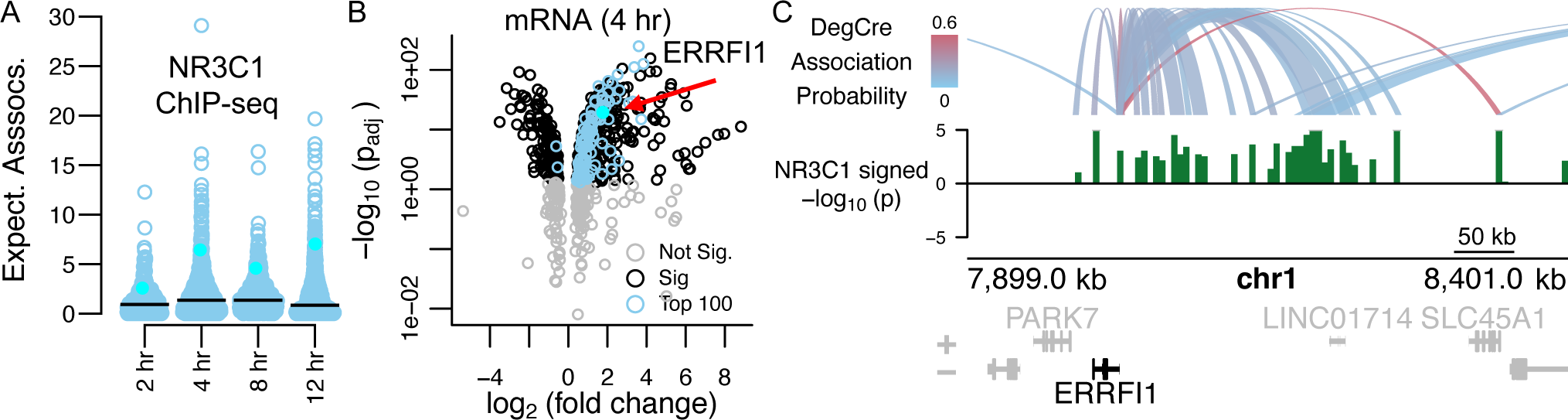
Identification of dexamethasone target genes with DegCre. A. The boxplot shows the distribution of expected DegCre associations per significant DEG (FDR less than or equal to 0.05) based on NRC31 ChIP-seq data from McDowell et al. The black line shows the median expected DegCre associations per DEG. The cyan points show values for *ERRFI1*. B. The volcano plot shows the −log_10_ of the adjusted (Bonferroni) differential expression p-value versus the log_2_ fold-change. Blue dots indicate genes whose expected number of associations is in the top 100 of all significant DEGs. C. The browser view shows DegCre associations (top panel) and NR3C1 ChIP-seq signal at four hours for an established glucocorticoid pathway target gene, *ERRFI1*. The NR3C1 signal is plotted as –log_10_ of the differential p-value multiplied by the sign of the log fold change. Regions of NR3C1 signal have been merged in some cases for better visibility at browser scale.

We further considered the DegCre results derived from NR3C1 (glucocorticoid receptor) ChIP-seq in McDowell et al. In these experiments, cells were treated with dexamethasone, a potent agonist of NR3C1. We looked at the subset of DEGs at four hours post treatment that had the top 100 highest expected associations with NR3C1 CREs (Figure 5B). This subset differs from what one would obtain by ranking DEGs by differential expression significance or fold-change magnitude alone, indicating that the DegCre-derived expected CRE associations per DEG adds additional, complementary information. *ERRFI1*, a well-established and ubiquitous target of NR3C1 (Juszczak and Stankiewicz 2018), exhibits significant but comparatively moderate fold-change (3.4-fold) in expression after NR3C1 stimulation (Figure 5B). However, *ERRFI1* ranks highly across all time points by expected associations per DEG (Figure 5A) due to the numerous regions of concordant differential NR3C1 occupancy associated to its expression by DegCre (Figure 5C).

## Discussion

DegCre represents a valuable advancement towards the fundamental goal of associating regulatory regions with gene expression. DegCre uses differential effects between conditions to identify regulatory regions specific to the causative perturbation. The probabilistic associations produced by DegCre span a wide range of interaction distances and enable the implementation of new analyses.

DegCre makes no parametric assumptions about the distribution of the input CRE and DEG p-values and thus has flexibility to accommodate a wide variety of data types. We noticed that the CRE p-values we produced from csaw analysis deviated somewhat from a uniform distribution, with high degree of “one” inflation (i.e., regions that show no differential signal). Because DegCre essentially uses the rank of these values, such unexpected distributions are not problematic. DegCre does not threshold by association distance or CRE p-value, avoiding issues that may arise from such practice.

We evaluated the validity of the DegCre associations by comparison with two orthogonal data types. First, we compared DegCre associations to HiC loops derived from the same cells (Figure 3). We initially conceived DegCre to primarily perform well for CREs proximal to DEG TSSs (e.g. within tens of kb). However, our comparison to the HiC loop calls indicated that DegCre can produce high-quality associations at distances at least up to 1Mb, particularly at early time points which are likely enriched for primary, direct CRE-DEG effects (Figure 3C and D). However, in cases where a CRE is near the TSSs of two significant DEGs, DegCre will generally assign it to the most proximal TSS, at times in conflict with HiC loops (Figure 3B). However, because HiC loops spanning short distances are rarer (Supplemental Figure 8A), the shorter DegCre association in Figure 3B may still be valid even if the failure to call the longer HiC-inferred association is likely a false negative (i.e., false negative loop calls, especially at shorter distances, are a possible explanation for such discrepancies). As such, we believe DegCre provides valuable, complementary information to HiC and related assays.

We also compared DegCre to CRISPR validated associations from Nasser et al. The authors compiled and generated this resource to benchmark various enhancer-to-promoter association methods, including their own, ABC (Nasser et al. 2021). As such, they were not specifically selected to be relevant to the perturbations and cell types used in our demonstration of DegCre. Very few of these CRISPR-tested associations overlapped with DegCre associations derived from the McDowell et al. data set (Supplemental File 3). DegCre associations from Savic et al. had a greater overlap but yielded moderate association probabilities for the overlapping regions (Supplemental Figure 10) but still showed clear separation of DegCre scores between validated and not validated CREs. DegCre generated many high-probability associations for other associations in both of these datasets, suggesting that the perturbations in these experiments altered gene expression primarily through CREs not tested by CRISPR in the Nasser et al. data set. However, the DegCre associations generated from Reed et al. had many overlaps with the CRISPR tested regions and yielded a large dynamic range of DegCre association probabilities, showing excellent agreement with CRISPR results (Figure 4, Supplemental Figure 10). Furthermore, the Reed et al. DegCre associations show comparable agreement with CRISPR validation as ABC scores (Figure 4, Supplemental Figure 10). The creators of ABC have demonstrated its superior performance to most other approaches for connecting enhancers to promoters in static (i.e., non-perturbation) experimental contexts (Nasser et al. 2021). The similar performance of DegCre to ABC in areas of predictive overlap thus strengthens the validity of DegCre associations.

Methods that rely on HiC or related techniques, such as ABC, lack the ability to identify shorter TSS to CRE associations due to the inherent difficulty in distinguishing such loops from background. DegCre operates well in this distance range. Additionally, implementation of HiC remains resource intensive, prohibiting its use in a variety of cell types and conditions. The resolution of HiC loop calls (size of the anchor regions) also depends on the sequencing depth (Rowley et al. 2020), such that high-resolution loop calls require even further resource expenditure. DegCre accepts data from less resource-intensive techniques, such as RNA-seq, ChIP-seq, and ATAC-seq, and returns at a resolution derived from those assays, generally at 100s of bp. We implemented csaw on relatively small bins to allow for subsequent merging to arrive at these feature sizes. Methods such as ABC seek to catalog all potential CRE to gene interactions in a single cell type and set of conditions, creating a valuable resource. We designed DegCre to place probabilities on subsets of those associations that depend on a given perturbation, viewing the system in a differential context.

We employ a penalty to CREs with multiple associations that favors the most proximal (Methods). This choice is supported by observations by Nasser et al. showing that a null model in which CREs are simply assigned to the closest TSS performs reasonably well (Nasser et al. 2021). Also, we believe this high performance is likely increased at the relatively short CRE to TSS distances observed for many high confidence DegCre associations (Figure 2A-C). However, there are clearly examples of a single CRE regulating several distal genes, such as the β-globin LCR (Grosveld et al. 1987). DegCre will likely underpredict associations for such regions, a limitation of this approach.

With default settings, DegCre requires concordance between CRE and DEG effect directions (both up or both down), although this requirement can be removed. For repressors, a more suitable requirement might be requiring opposite directions. Further, for CRE signals that are thought to lead to both directions within a given experiment (i.e., measurement of a TF that represses some targets and activates others), this assumption may best be ignored. We have not analyzed datasets under these alternative scenarios.

In its current implementation, DegCre accepts only one type of CRE signal as an input; consideration of multiple CRE inputs requires multiple independent runs of DegCre. We ultimately envision an implementation which accepts multiple CRE signals simultaneously and produces a composite association probability. Different CRE inputs are likely to have varying degrees of correlation, making their integration more challenging. Also, different CRE inputs will likely occur at non-overlapping genomic regions in some cases, further complicating the amalgamation of the signal. We are currently working on ways to overcome these challenges and create a multivariate version of DegCre.

To facilitate a clear presentation, we chose to use p-values derived from differential expression or differential CRE signal analyses using simple models. For example, we compared time zero separately to each successive time point. However, both csaw and DESeq2, and other similar methods, can use more complex regression models involving continuous variables and several covariates. We anticipate the application of DegCre to experimental designs warranting more complex regression analyses, such as drug treatment of a panel of patient-derived cell lines in which numerous covariates would require consideration.

We have presented DegCre, an algorithm for the probabilistic association of CREs with DEGs in response to perturbations. A freely available R package, DegCre produces convenient data structures and runs efficiently on large data inputs. We demonstrated its application to three distinct data sets, each involving different perturbations, cell contexts, and regulatory signal measurements. From these, DegCre produced associations involving established regulatory regions that confirm by two orthogonal methods, yielding associations that identify direct target genes of the perturbations. DegCre complements existing approaches by providing probabilistic scores for CRE-to-gene associations at a wide range of biologically relevant distances, using less-resource intensive and thus more broadly obtainable input data. We believe that DegCre is an important tool for the systematic and quantitative characterization of differential gene regulation.

## Methods

### DegCre package and algorithm

DegCre is an R package that is freely available on GitHub (https://github.com/brianSroberts/DegCre). (We are preparing to submit to Bioconductor). Documentation of the included functions is provided in the package manual. It operates within the GenomicRanges (Lawrence et al. 2013) framework, accepting GRanges objects as inputs and returning a Hits object with results as metadata. It includes functions for secondary calculations, visualization of results, and conversion to other formats. We ran DegCre using R version 4.2.1.

DegCre uses functionality within the GenomicRanges package to create overlaps and the associated distances between supplied TSS and CRE GRanges inputs. Next, DegCre bins the associations by TSS to CRE distance. The bin containing the longest associations is larger than the other bins to accommodate the remainder that occurs when the total number of associations is not an integer multiple of the number of bins. DegCre next attempts to balance high resolution (many bins with fewer associations) versus the minimization of the per bin CRE p-value distribution deviation from the global (un-binned) distribution. For an array of potential bin sizes (number of associations per bin), DegCre calculates the median Kolmogorov-Smirnov (KS) test statistic across all binned CRE p-value distributions versus the global distribution. DegCre picks the smallest bin size (containing the fewest associations per bin) that is less than a user specified fraction (defaults to 0.2) of the range from the lowest to highest median KS test statistic (Supplemental Figure 12). We chose this fraction threshold because it often occurred near an inflection point in the curve (Supplemental Figure 12).

To calculate the raw association probability for a given association between DEG i and CRE j, a_raw,i,j_, DegCre considers A_i,j_, the set of associations within the same distance bin that have CRE p-values as or more significant than p_CRE,j_ (Figure 1A,D). For this set, using Bonferroni corrected p-values, the expected number of true DEGs, E_DEG_, is equal to:

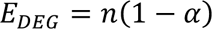

where α equals the chosen significance threshold of the adjusted DEG p-values, and n equals the number passing the threshold (Finner and Roters 2002). To generate a_raw,i,j_, one divides E_DEG_ by the size of set A_i,j_:

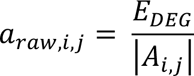

This probability is the average probability of association with a true DEG across the A_i,j_. For the considered association itself, we obtain a corrected probability a_corr,ij_, by setting the probability at zero if the adjusted DEG p-value is greater than α, since it cannot be an association between a CRE and a “true DEG” (Figure 1E). If the adjusted DEG p-value is less than or equal to α, no correction is made.

A given CRE will often have associations to multiple genes (Figure 1F). We favor the more proximal associations by down-weighting more distal associations in proportion to the sum of the more proximal. The final association probability a, is derived by:

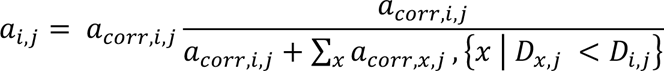

where D is the maximum association distance of the bin to which a given association belongs. Associations from the same CRE to different DEGs in the same distance bin will not penalize each other. Associations in the first (shortest) distance bin will never be altered by this process. The process moves in order of increasing distance bins, such that association probabilities are down-weighted by more proximal association probabilities that have been down-weighted already.

DegCre calculates a type of false discovery rate (FDR) for association probabilities. As a null hypothesis, a_bin_, we consider the true DEG association probability of a given distance bin without regard to the ordering of association CRE p-values (Figure 1D). If all associations in the bin had the same CRE p-value their association probabilities would be a_bin_. As described above, the process of calculating a_i,j_ involves considering the distance bin subset A_i,j_ and calculating E_DEG_, the expected number of true DEGs. This process can be modeled as a set of Bernoulli trials in which the number of trials is the size of A_i,j_, or |A_i,j_ |, the number of successes is E_DEG_, and the probability of success is a_bin_. The FDR of the association in this case is the probability that a given association probability exceeds the value derived its association distance alone, and is given by:

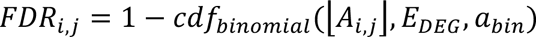

### Data processing and visualizations

We downloaded the presented data sets from public repositories. All accession numbers are provided in Supplemental File 1. For ChIP-seq and ATAC-seq data from Reed et al. and Savic et al. we aligned the FASTQ files to hg38 using bowtie2 version 2.3.5.1 (Langmead and Salzberg 2012) and processed with samtools version 1.16.1 (Danecek et al. 2021) to bam files. We obtained bam files directly for McDowell et al. data. We derived log-fold changes and p-values associated with GRanges from the bam files using the R package csaw version 1.32.0 (Lun and Smyth 2016) in R version 4.2.1. For csaw analysis of TF ChIP-seq data, we used 20 bp windows and kept the top 0.5% with highest signal for differential analysis. For open chromatin assays, histone ChIP-seq, and RNA Pol2 ChIP-seq, we used 200 bp windows and kept the top 2% with highest signal. We selected these values by finding those that produced the best clustering of samples by treatment and time point within experiment sets. Within csaw, we made comparisons to the zero time or control conditions for each time point. We obtained gene count tables for all RNA-seq data. We calculated log-fold changes and p-values for each time point relative to the zero time point or control using the R package DESeq2 version 1.38.3 (Love et al. 2014). We associated these values with all TSSs for each gene using annotations from EPDNew (Dreos et al. 2015; Meylan et al. 2020), yielding GRanges. We lifted over HiC loop calls from Reed et al. to hg38 using the R package rtracklayer version 1.58.0 (Lawrence et al. 2009). We obtained CRISPR experimental data and associated ABC calls from Supplemental Table 5 from Nasser et al. (Nasser et al. 2021). We lifted over this data to hg38 using rtracklayer version 1.58.0. We made all data visualizations using R version 4.2.1, in some cases using DegCre built-in functions. We made the browser plots using functions that use the R package plotgardener (Kramer et al. 2022).

## Acknowledgements

This work was supported by the Leo Fund. We thank the members of Myers Lab for useful discussions and suggestions. We thank Dr. E. Christopher Partridge for editing the manuscript.

## Data Access

We did not generate any previously unpublished data for this manuscript.

